# Highly prevalent cargo genes of F plasmids harbored in pathogenic *Escherichia coli*

**DOI:** 10.64898/2026.11.04.764203

**Authors:** S Guirao-Rico, SC Flament-Simon, C Madrid, C Balsalobre, M Hüttener, J.E. Blanco, J Blanco, J Rozas, A Juárez

**Author notes:** Both authors contributed equally to this work (S Guirao-Rico, SC Flament-Simon).

## Abstract

Analysis of prevalent orthogroups (OGs) in a collection of F plasmids from extraintestinal *Escherichia coli* ST131 isolates has led to the identification of two groups of previously uncharacterized cargo genes that may be widely distributed among F plasmids of pathogenic *E. coli*.

Group 1 comprises four OGs whose encoded proteins are predicted to be involved in transport-related functions. The corresponding genes cluster with other genes associated with metabolism and are highly prevalent in extraintestinal ST131 isolates.

Group 2 comprises several OGs whose encoded proteins currently have unknown functions. These OGs are present in a large proportion of F plasmids from both intestinal and extraintestinal *E. coli* isolates. The corresponding genes cluster within an F-plasmid region that also contains genes involved in DNA metabolism, suggesting that the proteins encoded by these newly identified OGs may likewise contribute to F-plasmid DNA biology.

Characterizing the functions of these newly identified proteins may improve our understanding of the contribution of F plasmid to *E. coli* virulence and reveal novel targets for antimicrobial intervention, either by limiting F-plasmid dissemination among multidrug-resistant (MDR) *E. coli* strains or by directly targeting plasmid-carrying bacteria.

## Introduction

Nowadays, antimicrobial resistance (AMR), often referred to as a “silent pandemic,” is recognized by the World Health Organization (WHO) as one of the greatest threats to public health (1) and it has increasingly become a concern beyond the scientific and medical communities. Despite the availability of antibiotics, bacterial infectious diseases remain the second leading cause of death worldwide (2, 3). In 2021, an estimated 4.71 million deaths were associated with AMR, meaning that resistance contributed to the burden of these infections. Of these, approximately 1.14 million deaths were directly attributable to AMR, indicating that they would likely not have occurred if the infecting bacteria had been susceptible to effective antibiotic treatment (4).

A major factor contributing to the spread of AMR is the ability of bacteria to transfer resistance determinants, even between distantly related organisms. Horizontal gene transfer (HGT) is a major mechanism driving the dissemination of AMR, with plasmids playing a pivotal role in this process. Plasmids facilitate the transmission of resistance determinants against all major classes of antimicrobials (5), particularly among Gram-negative bacteria (6–8), but also in Gram-positive bacteria (9). Bacterial plasmids are highly diverse and can be classified using different approaches. One of the most widely used and well-established classification systems is based on incompatibility (Inc) groups (10).

The central role of plasmids in the dissemination of AMR, together with the ability of bacteria to rapidly acquire resistance to newly introduced antibiotics, highlights the limitations of approaches based solely on antibiotic development, as resistance to newly introduced agents can rapidly emerge. Modifications of existing antibiotic classes have achieved limited success and do not prevent the eventual emergence of resistance. Moreover, only two completely new antibiotic classes have successfully reached clinical use over the past decades (11), highlighting the urgent need to identify novel antimicrobial targets and develop alternative therapeutic strategies.

These limitations have prompted the development of alternative approaches to combat AMR, including the so-called Eco-Evo strategies (12). These approaches are based on the recognition that AMR is widespread in natural microbial communities. Rather than attempting to kill MDR microorganisms, Eco-Evo strategies aim to prevent the emergence and spread of resistant bacteria in the environment or restore bacterial populations that remain susceptible to antibiotics. Examples of such approaches include the development of novel vaccines (13) and the inhibition of bacterial plasmid conjugation (14, 15).

Achieving these objectives requires a deeper understanding of plasmid biology. Although considerable progress has been made in elucidating key aspects of plasmid replication and conjugative transfer, many features of plasmid biology remain poorly understood. As with other genomic elements, sequencing of bacterial plasmids frequently reveals numerous open reading frames (ORFs) with unknown functions, which in some cases account for more than 50% of the predicted genes. Elucidating the functions of these genes could substantially improve our understanding of how bacterial plasmids contribute to host fitness and, consequently, to bacterial survival and persistence under challenging environmental conditions.

One example illustrating the potential of this approach has been the identification and characterization of bacterial immunoglobulin-like (Big) domain-containing proteins encoded by plasmids belonging to different incompatibility groups, particularly IncHI and IncA/C. These high-molecular-mass proteins associate with extracellular bacterial appendages involved in plasmid conjugative transfer (15, 16). Targeting these proteins has proven effective both in protecting mice against infection with a *Salmonella* strain carrying an IncHI resistance plasmid and in interfering with the conjugative transfer of these plasmids (17). IncHI plasmids are widely distributed among enterobacteria and are particularly prevalent in *Salmonella* isolates (18). IncA/C plasmids have likewise been identified in several enterobacterial isolates (19). This example illustrates how functional characterization of previously uncharacterized plasmid-encoded proteins can reveal both biological functions and potential targets for controlling AMR.

A particularly important group of plasmids in this context are F plasmids. Both *Escherichia coli* (*E. coli*) and *Klebsiella pneumoniae* (*K. pneumoniae*) strains producing extended-spectrum β-lactamases (ESBLs) have spread globally and represent a major public health concern (20). In MDR *E. coli* and *K. pneumoniae* isolates, F plasmids are frequently detected (19). In *E. coli*, F plasmids play a central role in the dissemination of multiple AMR determinants, including those encoding ESBLs and carbapenemases (19, 21, 22). These plasmids are important genetic elements in both commensal (23) and pathogenic *E. coli* isolates. Their widespread distribution among both commensal and pathogenic *E. coli* underscores their importance as genetic elements shaping the biology and evolution of this species (for a review, see ref. 24).

Comparative genomic studies have revealed a high degree of heterogeneity among F plasmids. These plasmids may carry different antimicrobial resistance determinants, as well as several virulence-associated functions, including the *ompP* gene, which encodes an outer membrane protease capable of degrading cationic antimicrobial peptides (25); determinants involved in iron acquisition and uptake (26); genes encoding heat-labile (LT) and heat-stable (ST) enterotoxins (27); and the predicted iron uptake system *cjrABC-senB* (28). This genetic diversity suggests that the biological roles of F plasmids extend beyond their established contributions to antimicrobial resistance and virulence. F plasmids may carry additional functions that contribute to bacterial fitness, persistence, or proliferation within the host. Identifying and characterizing these functions may therefore provide new insights into the biological roles of F plasmids.

Among pathogenic *E. coli*, extraintestinal pathogenic *E. coli* (ExPEC) are responsible for a wide range of severe infections, most commonly urinary tract infections (UTIs) and bloodstream infections, but also pneumonia, neonatal meningitis, and sepsis. ExPEC isolates can be classified into distinct sequence types (STs) based on multilocus sequence typing (MLST). Several major STs, including ST69, ST73, ST95, ST131 and ST1193, comprise ExPEC lineages associated with severe disease (29–31). Of these, ST131 is globally widespread and is characterized by a high prevalence of multidrug-resistant (MDR) isolates associated with UTIs and bloodstream infections (22, 29, 32–36). ST131 comprises several distinct evolutionary lineages, including clades A, B, and C, with further diversification of clade C into subclades C0, C1-M27, non-C1-M27, and C2 (H30Rx). The diversification of these lineages has been associated with the acquisition and evolution of mobile genetic elements, particularly IncF plasmids, which are considered important drivers of the global dissemination of ST131 (37). Despite their established association with the success of ST131, the specific contributions of many plasmid-encoded functions to the fitness, persistence, and pathogenicity of these lineages remain poorly understood.

Based on these previous considerations, the present study sought to identify highly prevalent, previously uncharacterized F-plasmid cargo genes whose products may contribute to the fitness of pathogenic *E. coli*. We initially analyzed a collection of 29 ST131 *E. coli* strains isolated and sequenced by members of our group, all of which harbored F plasmids. We reasoned that ORFs consistently present in these plasmids but not associated with established core F-plasmid functions, could represent conserved cargo genes with potential roles in bacterial fitness. We then assessed the distribution of these ORFs among the F plasmids carried by the ST131 isolates and their prevalence in a larger collection of F plasmids from other *E. coli* lineages.

Our analysis identified several previously uncharacterized F-plasmid-encoded genes that are highly conserved among F plasmids carried by extraintestinal ST131 *E. coli* isolates. Notably, some of the most prevalent F-plasmid genes identified in ST131 were also widely distributed among F plasmids from other pathogenic *E. coli* isolates, suggesting that these genes are not restricted to the ST131 lineage. Characterizing the functions of these conserved genes may provide new insights into the contribution of F plasmids to the fitness of pathogenic *E. coli* and may ultimately identify novel targets for strategies aimed at limiting F-plasmid dissemination or developing antimicrobial approaches against F-plasmid-carrying bacteria.

## Materials and Methods

### Epidemiological background of the *E. coli* collection

A collection of *Escherichia coli* ST131 clinical isolates was used in this study. Specifically, we analyzed the F plasmids of 29 extended-spectrum β-lactamase (ESBL)-producing extraintestinal pathogenic *Escherichia coli* (ExPEC) belonging to the Sequence Type Complex 131 or ST131 lineage. The isolates were recovered in 2015 from two European hospitals: the Hospital Universitario Lucus Augusti in Lugo (Spain, *n* = 26) and the Beaujon Hospital in Paris (France, *n* = 3). Regarding their clinical origin, most of the strains were isolated from urine samples (*n* = 27), while the remaining two isolates were recovered from blood (*n* = 1) and bone tissue (*n* = 1) (38; Supplementary Table S1).

### DNA extraction, whole-genome sequencing, assembly and annotation

Total DNA was extracted with the QIAmp DNA Mini Kit (Qiagen). Libraries were prepared using the TruSeq DNA PCR-Free protocol (Illumina) at the Genomics and Bioinformatics Core Facility (Centre for Biomedical Research of La Rioja, CIBIR). Paired-end 100 bp reads were generated on an Illumina HiSeq 1500 from libraries with an average insert size of 550 bp. Genomes were assembled at the Genomics and Bioinformatics Core Facility (Centre for Biomedical Research of La Rioja, CIBIR) and reconstructed using PLACNET webserver (39). The assembly information of draft genomes can be found in Supplementary Table S2. Identification of Open Reading Frames (ORFs) and genome annotation of the assembled genetic elements was performed using Prokka (40). Genomes were *in silico* typed following the methodology described in the Supplementary Methods. The datasets for the 29 ESBL-producing ST131 genomes of human origin can be found in the NCBI sequence databases as part of BioProject PRJNA843680: Genomic characterization of MDR resistant ESBL *Escherichia coli* strains from human origin, accessions for this submission are shown in Supplementary Table S1.

### Orthology inference and functional annotation of orthologous groups in 29 ST131 strains

Orthology relationships among the gene products encoded in the IncF plasmids of 29 extraintestinal E. coli (ExPEC) strains belonging to sequence type complex 131 (ST131) were identified using Broccoli v1.2 (41) (Supplementary Tables S3 and S4). For each orthology group (hereafter called OG), functional annotation was performed using three different approaches: i) running BLASTP, using BLAST+ v2.16.0 (42, 43), against the SwissProt database (release-2024_04) restricted to bacterial sequences; ii) running eggnog-mapper v2.1.10 (44, 45) and iii) running InterProScan v5.57 (46) with InterPro database release 90.0 (release July 2022) to have additional information regarding the putative function of each orthogroup. For each approach, searches were performed using as query either i) the longest protein of each orthogroup or ii) all the proteins of each orthogroup. From the initial set of orthogroups (OGs), we selected only those (13 OGs) present in a high percentage of the 29 bacterial strains (> 80%) with significant BLASTP *E*-values (< 1 × 10⁻^9^) (see Results section).

### COMPASS database analysis

To investigate the distribution of selected orthologous groups (OGs) among IncF plasmids, we used the COMPASS database, downloaded from https://github.com/itsmeludo/COMPASS/tree/master, a curated collection of 12,084 complete plasmids and associated metadata representing 1,571 bacterial species sampled worldwide over more than a century (47). Using the metadata provided in the COMPASS database, plasmid sequences corresponding to *E. coli* IncF plasmids were selected, yielding a filtered dataset of 578 plasmids (Supplementary Table S5).

The longest protein sequence from each selected OG was used as a query in TBLASTN searches (BLAST+ v2.16.0; 42, 43) against each plasmid in the filtered COMPASS dataset, using an *E*-value of 1 × 10⁻⁵. In addition, for each plasmid, the number and proportion of selected OGs producing significant matches were determined relative to the total number of selected OGs analyzed.

Metadata associated with the filtered plasmid dataset were used to investigate plasmid characteristics, including host pathogenicity status and host isolation source when available. The metadata were retrieved programmatically from the NCBI databases using the Entrez utilities implemented in Biopython v1.84 under Python v3.9. Custom Python scripts were used to parse and process the retrieved records. Isolation source information was available for 112 of the 578 IncF plasmids (19.4%).

### Reid *et al*. dataset analysis

We used the strain collection derived from the *E. coli* bloodstream infection (BSI) dataset described by Reid et al. (48), comprising 4,711 isolates collected in the United Kingdom (Cambridge and Oxford hospitals; 2001-2018), France (multiple hospitals; 2005 and 2016-2017), and Australia (a single hospital; 2013-2016). Reid et al. compiled this dataset from four previously published studies whose authors had already applied sampling strategies to minimize the inclusion of highly similar or identical isolates; consequently, no additional filtering was performed by Reid et al. for this purpose.

Publicly available genomic data were retrieved from NCBI using the NCBI Datasets command-line interface (ncbi-datasets-cli v18.5.0) for assembly accessions (GCA/GCF) and NCBI Entrez Direct (EDirect v24.0; esearch and efetch) for nucleotide and WGS contig accessions. Although Reid et al. identified 3,463 isolates carrying at least one IncF replicon sequence typing (RST) allele, many of the corresponding genome assemblies (all of them from Oxford) were not publicly available and only raw sequencing reads could be accessed. While Reid et al. generated assemblies for these isolates, the resulting assemblies were not deposited in public repositories and were therefore unavailable for this study. Consequently, only isolates with publicly available assemblies or contigs (i.e. those from France, Australia and Cambridge) were retained for downstream analyses.

We used the F+ dataset comprising 602 isolates carrying at least one IncF Replicon Sequence Type (RST) allele.

The longest protein of each selected OG was searched against the corresponding nucleotide databases (i.e. multifasta database comprising the 602 isolates) using TBLASTN (BLAST+ v2.16.0; 42, 43) with an *E*-value of 1 × 10⁻⁵. For each selected OG, the number and proportion of isolates with and without significant matches were calculated overall and stratified by sampling origin (France, Australia and Cambridge).

### Stephens *et al*. dataset analysis

To evaluate the distribution of selected OGs in commensal E. coli, we analyzed the complete-genome collection described by Stephens et al. (23). Of the 50 strains included in the original dataset, four lacked plasmid sequence information and were excluded, resulting in a final dataset of 46 strains comprising 156 plasmids. Plasmid accession numbers and associated metadata were retrieved from Supplementary Table S1 of Stephens et al. (23), and plasmid nucleotide sequences were downloaded from NCBI.

All plasmid sequences were combined into a multifasta database. The longest protein of each selected OG was used as query in TBLASTN (BLAST+ v2.16.0; 42, 43) searches against this plasmid database (*E*-value ≤ 10^-5). For each OG, we determined the number and proportion of plasmids and strains producing significant hits. Associations between OG occurrence and plasmid- or strain-level characteristics were assessed, including antimicrobial resistance, multidrug resistance, IncF replicon content, plasmid size category, and host phylogroup composition.

Enrichment analyses were performed using hypergeometric tests to determine whether specific characteristics (e.g., multidrug resistance, resistance-associated plasmids, or IncF carriage) were overrepresented among plasmids or strains producing significant hits relative to their frequency in the complete dataset.

### Association of candidate orthogroups with functionally annotated F plasmid proteins

To associate the identified OGs with functionally annotated proteins, BLAST searches using BLAST+ v2.16.0 (42, 43) were performed using the annotated protein sequences of two F plasmids from the ST131 strain collection as queries. The plasmids analyzed were pEIC35 and pEIC39, isolated from the ST131 strains LREC268, and LREC291, respectively. The protein sequences encoded by each plasmid were searched independently against a database containing all protein sequences from all identified OGs using BLASTP with an *E*-value threshold of 1 × 10⁻⁵. Significant matches were used to identify OGs corresponding to annotated proteins of the functionally characterized F plasmids.

### Comparative plasmid genome and visualization

Protein annotation files for pEIC35 and pEIC39 plasmids, together with their corresponding nucleotide FASTA sequences of the F plasmid, were used to generate GenBank files using an in-house Python script (Python v.3.14.6) based on Biopython v.1.87 (49) and pandas v3.0.5 package. The script combined nucleotide sequences with the corresponding protein annotation coordinates to produce annotated GenBank files suitable for comparative genomic visualization. The resulting GenBank files were used to compare the two using the pgv-blast workflow implemented in pyGenomeViz v. 1.7.0. (50; Available from: https://github.com/moshi4/pyGenomeViz). Protein-level sequence similarities were identified using the following parameters: --seqtype protein, --identity_thr 30, and --length_thr 100. Comparative genomic maps were generated with a scale bar (--show_scale_bar), curved links between homologous regions (--curve), gene labels (--feature_labeltype gene), a label size of 5 (--feature_labelsize 5), and exported in PNG, HTML, and SVG formats (--formats png html svg).

## Results

### Plasmidome of the ST131 strains

Reconstruction using PLACNET allowed us to characterize a presumptive plasmidome for each genome. In total, 86 plasmids were identified across the 29 genomes comprising the ST131 group (Supplementary Table S6). Comparative analysis of these plasmidomes revealed several common features associated with the different ST131 clades, which are summarized in Supplementary Tables S6 and S7. The bioinformatic analysis indicated that these strains carried diverse ESBL-encoding *bla* genes and a total of 31 IncF plasmids (Supplementary Tables S3 and S7). Despite the considerable plasmid diversity observed among ESBL-producing ST131 *E. coli* isolates, most of the identified plasmids were assigned to the MOBF, MOBP, and MOBQ relaxase families, which together represented the major component of the plasmidome. In addition, Col-like and IncF plasmids were the most prevalent. As expected, most virulence- and antimicrobial resistance-associated genes were located on IncF plasmids.

Determination of the highly prevalent orthogroups in the F plasmids of 29 extraintestinal ST131 *E. coli* strains.

We next sought to identify highly prevalent genes in the F plasmids of the analyzed ST131 strains and to determine whether, in addition to core F-plasmid genes, previously uncharacterized cargo genes were conserved among these plasmids.

We inferred the orthology relationships among genes identified in their corresponding F plasmids. The resulting orthogroups (OGs) are listed in the Supplementary Tables S4 and S8. For further studies, we selected those OGs whose genes were present in at least 83% of the sequenced F plasmids (Table 1). OGs corresponding to proteins involved in core plasmid functions such as plasmid replication and conjugative transfer, as well as transposases and antibiotic resistance determinants, were excluded from the analysis (e.g., OGs 482, 511, 3, 18, 838, 321).

**Table 1.** Most prevalent orthogroups (OGs) identified in F plasmids across the ST131 strain collection. The 13 OGs selected for further analysis are shown in bold.

| **Orthogroup (OG) ID** | **Best BLAST hit annotation** |
|---|---|
| **OG 495** | **Putative antirestriction protein YubI** |
| **OG 44** | **Toxin CcdB** |
| **OG 329** | **Uncharacterized protein YubA** |
| **OG 336** | **Uncharacterized protein YubJ** |
| **OG 337** | **Uncharacterized protein YubJ** |
| **OG 338** | **Uncharacterized protein YubG** |
| **OG 339** | **Uncharacterized protein YubG** |
| OG 482 | Replication initiation protein |
| OG 497 | Putative methylase YubD |
| OG 511 | Replication regulatory protein repA2 |
| OG 43 | Antitoxin CcdA |
| OG 46 | Ribosomal RNA small subunit methyltransferase A |
| **OG 73** | **Protein SopA** |
| OG 496 | Uncharacterized protein YubE |
| OG 3 | Plasmid-derived single-stranded DNA-binding protein |
| OG 18 | Transposase for insertion sequence-like element IS431mec |
| OG 38 | Coupling protein TraD |
| OG 47 | Ribosomal RNA small subunit methyltransferase A |
| OG 49 | - |
| OG 77 | Uncharacterized ABC transporter ATP-binding protein YvrO |
| OG 321 | Beta-lactamase CTX-M-14 |
| **OG 335** | **UPF0380 protein YubP** |
| OG 422 | Uncharacterized 24.3 kDa protein |
| OG 498 | Resolvase |
| OG 506 | Protein PsiB |
| OG 507 | Protein PsiA |
| OG 9 | Multifunctional conjugation protein TraI |
| OG 36 | Multifunctional conjugation protein TraI |
| OG 48 | - |
| OG 75 | Uncharacterized protein Lin2408 |
| **OG 76** | **Pyoverdine export ATP-binding/permease protein PvdT** |
| **OG 78** | **Iron permease EfeU** |
| **OG 131** | **Macrolide export ATP-binding/permease protein MacB** |
| **OG 421** | **UPF0423 protein BMEII0885** |
| OG 480 | Antitoxin PemI |
| OG 504 | UPF0401 protein YubL |

We retained three categories of highly prevalent OGs: (i) the most prevalent, OG495, present in 97% of the strains and encoding a putative antirestriction protein; (ii) six OGs with a prevalence of 93 to 86% that encoded proteins with unknown function (OGs 329, 336, 337, 338, 339, 335), and (iii) four OGs encoding proteins predicted to be involved in membrane trafficking (OGs 76, 78, 131, 421). The lengths of the corresponding protein sequences are provided in Supplementary Table S9. For subsequent analyses, we used these three groups of OGs, together with two OGs associated with established plasmid functions -OG 44, encoding the CcdB toxin, and OG 73, encoding the SopA protein- which were included as controls for assessing the prevalence of conserved core plasmid genes across the different F-plasmid collections.

### Presence of the selected orthogroups in the F plasmids of *E. coli* strains in the COMPASS database

We next assessed whether the most prevalent OGs identified in the ST131 strains were also present in F plasmids from other *E. coli* strains. To this end, the selected OGs were used as queries against the 578 *E. coli* F-plasmid sequences extracted from the COMPASS database (47), which comprises a large collection of plasmids from *E. coli* strains, including predominantly pathogenic isolates.

The results revealed several distinct patterns of OG distribution (Table 2; Supplementary Table S5). Fifty-two strains (9% of total strains) contained 100% of the selected OGs. Sixty-five strains (11.2%) contained 92% of the OGs, typically lacking SopA. One hundred and nine strains (18.8%) contained 69% of the OGs. In these strains, all four OGs related to membrane trafficking were absent. Two hundred one strains (34.7%) contained 61% of the OGs. In addition to the OGs absent in the preceding group, OGs 73, 329 or 44 were absent in some strains. Three hundred and seven strains (53.11%) retained 53% of the OGs, with OGs 336, 337, 338, 339 and 495 present in all these strains, whereas OGs 73, 329, 335 or 44 showed variable patterns of presence and absence. Four hundred and twenty strains (72.6%) retained 46% of the OGs, displaying a distribution pattern similar to that of the preceding group, with OGs 337, 495, 336, 335, 338 and 339 present. Finally, 461 strains (79.7%) retained only 30% of the selected OGs, with OGs 338 and 339 remaining frequently present.

**Table 2.** BLAST results for 13 selected orthogroups (OGs) against a filtered COMPASS database restricted to 578 F-plasmid sequences from *Escherichia coli*.

| Orthogroup (OG) ID | Number of plasmids with a BLAST hit | Percentage of plasmids with a BLAST hit | Best BLAST hit | Percentage of ST131 strains with a BLAST hit |
|---|---|---|---|---|
| OG 495 | 436 | 75 | Putative antirestriction protein YubI | 97 |
| OG 44 | 254 | 43 | Toxin CcdB | 93 |
| OG 329 | 323 | 55 | Uncharacterized protein YubA | 93 |
| OG 336 | 433 | 74 | Uncharacterized protein YubJ | 93 |
| OG 337 | 440 | 76 | Uncharacterized protein YubJ | 93 |
| OG 338 | 450 | 77 | Uncharacterized protein YubG | 93 |
| OG 339 | 450 | 77 | Uncharacterized protein YubG | 93 |
| OG 73 | 288 | 49 | Protein SopA | 90 |
| OG 335 | 383 | 66 | UPF0380 protein YubP | 86 |
| OG 76 | 71 | 12 | Pyoverdine export ATP-binding/permease protein PvdT | 83 |
| OG 78 | 72 | 12 | Iron permease EfeU | 83 |
| OG 131 | 72 | 12 | Macrolide export ATP-binding/permease protein MacB | 83 |
| OG 421 | 72 | 12 | UPF0423 protein BMEII0885 | 83 |

These data indicate that OGs 76, 78, 131 and 421, which were among the most prevalent OGs in the F plasmids of the ST131 strains analyzed, are not widely distributed among the F plasmids represented in the COMPASS collection, being present in only approximately 11% of the plasmids analyzed. In contrast, OG495 and OGs 329, 335, 336, 337, 338 and 339 were widely distributed across the F plasmids in this collection. The latter OGs encode proteins of unknown function (Yub proteins). Several of these OGs were present in nearly 80% of the F plasmids analyzed.

We next examined whether the distribution of the selected OGs was associated with the reported source of the *E. coli* strains. Despite incomplete metadata regarding the isolation source of some strains, a trend was apparent (Supplementary Table S5). F plasmids from strains of extraintestinal origin (i.e., isolates recovered from urine, blood, or cases of septicemia) contained 69–100% of the analyzed OGs, including those associated with membrane-trafficking functions (Supplementary Table S5). Strains associated with intestinal infections (e.g., isolates recovered from feces, rectal swabs, or patients with diarrhea) generally harbored F plasmids containing up to 69% of the analyzed OGs, although plasmids containing 61% or fewer of the OGs were more frequent. These plasmids generally lacked the OGs associated with membrane-trafficking functions.

To further support the hypothesis that OGs 76, 78, 131 and 421 are predominantly associated with strains causing extraintestinal infections, we performed an additional BLAST analysis against 602 extraintestinal *E. coli* strains from Reid *et al*. dataset (48). OGs 76, 78, 131 and 421 were detected in approximately 50% of the strains analyzed (Table 3), whereas OGs encoding the Yub proteins were detected in more than 70% of the strains (Table 3).

**Table 3.** Blast results for the 13 selected orthogroups (OGs) against a collection of 602 pathogenic extraintestinal *Escherichia coli* strains.

| Orthogroup (OG) ID | Best BLAST hit | Number of strains with no BLAST hit | Number of strains with a BLAST hit | Non-blasted/Total (Ratio) | Blasted/Total (Ratio) |
|---|---|---|---|---|---|
| OG 495 | Putative antirestriction protein YubI | 175 | 427 | 0.29 | 0.71 |
| OG 44 | Toxin CcdB | 198 | 404 | 0.33 | 0.67 |
| OG 329 | Uncharacterized protein YubA | 182 | 420 | 0.30 | 0.70 |
| OG 336 | Uncharacterized protein YubJ | 169 | 433 | 0.28 | 0.72 |
| OG 337 | Uncharacterized protein YubJ | 177 | 425 | 0.29 | 0.71 |
| OG 338 | Uncharacterized protein YubG | 171 | 431 | 0.28 | 0.72 |
| OG 339 | Uncharacterized protein YubG | 171 | 431 | 0.28 | 0.72 |
| OG 73 | Protein SopA | 200 | 402 | 0.33 | 0.67 |
| OG 335 | UPF0380 protein YubP | 117 | 485 | 0.19 | 0.81 |
| OG 76 | Pyoverdine export ATP-binding/permease protein PvdT | 321 | 281 | 0.53 | 0.47 |
| OG 78 | Iron permease EfeU | 323 | 279 | 0.54 | 0.46 |
| OG 131 | Macrolide export ATP-binding/permease protein MacB | 322 | 280 | 0.53 | 0.47 |
| OG 421 | UPF0423 protein BMEII0885 | 323 | 279 | 0.54 | 0.46 |

### Presence of the selected orthogroups in the F plasmids of commensal *E. coli* strains

The above reported data correlate the identified OGs with F plasmids from pathogenic *E. coli* strains. To complete our analysis, we also investigated whether the selected OGs are present in F plasmids from *E. coli* strains considered to be commensal. For this analysis, we used the data from the study of Stephens *et al*. 2020 (23), which generated genomic sequence data for 156 plasmids harbored by 46 commensal *E. coli* strains.

Analysis of the presence and absence of the selected OGs in the individual plasmids (Table 4; Supplementary Table S10), revealed a markedly lower prevalence than that observed in the F plasmids of either the ST131 strains or the *E. coli* strains represented in the COMPASS database. The group of OGs encoding proteins associated with membrane trafficking was detected in 14% of the plasmids. The prevalence of the remaining selected OGs ranged from 16% to 32% of the plasmids analyzed. The prevalence of the remaining selected OGs ranged from 16% to 32% of the plasmids analyzed.

**Table 4.** BLAST results for 13 selected orthogroups (OGs) against the complete plasmidome of each strain of a commensal *Escherichia coli* collection consisting in 46 strains and 156 plasmids.

| Orthogroup (OG) ID | Number of plasmids with a BLAST hit | Percentage of plasmids with a BLAST hit | Best BLAST hit | Number of strains with a BLAST hit | Percentage of strains with a BLAST hit |
|---|---|---|---|---|---|
| OG 495 | 51 | 32 | Putative antirestriction protein YubI | 39 | 84 |
| OG 44 | 30 | 19 | Toxin CcdB | 30 | 65 |
| OG 329 | 44 | 28 | Uncharacterized protein YubA | 33 | 71 |
| OG 336 | 52 | 33 | Uncharacterized protein YubJ | 40 | 86 |
| OG 337 | 52 | 33 | Uncharacterized protein YubJ | 40 | 86 |
| OG 338 | 51 | 32 | Uncharacterized protein YubG | 39 | 84 |
| OG 339 | 51 | 32 | Uncharacterized protein YubG | 39 | 84 |
| OG 73 | 25 | 16 | Protein SopA | 24 | 52 |
| OG 335 | 36 | 23 | UPF0380 protein YubP | 33 | 71 |
| OG 76 | 22 | 14 | <sup>a</sup>Pyoverdine export ATP-binding/permease protein PvdT | 22 | 47 |
| OG 78 | 22 | 14 | <sup>b</sup>Iron permease EfeU | 22 | 47 |
| OG 131 | 22 | 14 | <sup>c</sup>Macrolide export ATP-binding/permease protein MacB | 22 | 47 |
| OG 421 | 22 | 14 | <sup>d</sup>UPF0423 protein BMEII0885 | 22 | 47 |

A different picture emerged when the analysis was extended from individual plasmids to the complete plasmidome of each commensal strain (Table 4, Supplementary Table S11). In this case, the selected OGs were detected in 47–86% of the 46 commensal strains analyzed. This difference is important because most of these strains harbor more than one plasmid, indicating that OGs that occur at relatively low frequencies in individual plasmids may nevertheless be widely represented within the overall plasmid content of commensal *E. coli* strains.

### Location of the genes corresponding to the identified orthogroups in the functional maps of F plasmids from ST131 strains

To determine the genomic location of the selected OGs and correlate them with functionally annotated proteins, we mapped the consensus sequence of each selected OG onto the functionally annotated sequences of two F plasmids from the ST131 strain collection analyzed in this study, pEIC35 and pEIC39 (GenBank pZ167158 and pZ167159 respectively)

These plasmids were isolated from ST131 strains LREC268 and LREC291, respectively. The functional annotations and corresponding OG assignments are provided in Supplementary Tables S12 and S13.

Examination of the functional genomic maps of pEIC35 and pEIC39 revealed two distinct patterns of organization (Figs. 1 and 2). The genes corresponding to OGs 78, 131, 76, and 421 were clustered within a specific genomic region in both plasmids. In contrast, the genes corresponding to OGs 329, 335, 336, 337, 338, and 339 formed a separate cluster located in a different region of the plasmids. We refer to these two regions hereafter as region T, named for the association of the OGs with transport-related functions and region Y, named for Yub proteins, respectively. Notably, the region containing the genes of OGs 76, 78, 131, and 421 showed a higher degree of conservation between pEIC35 and pEIC39 than the region containing the Yub-associated OGs.

**Figure 1.**
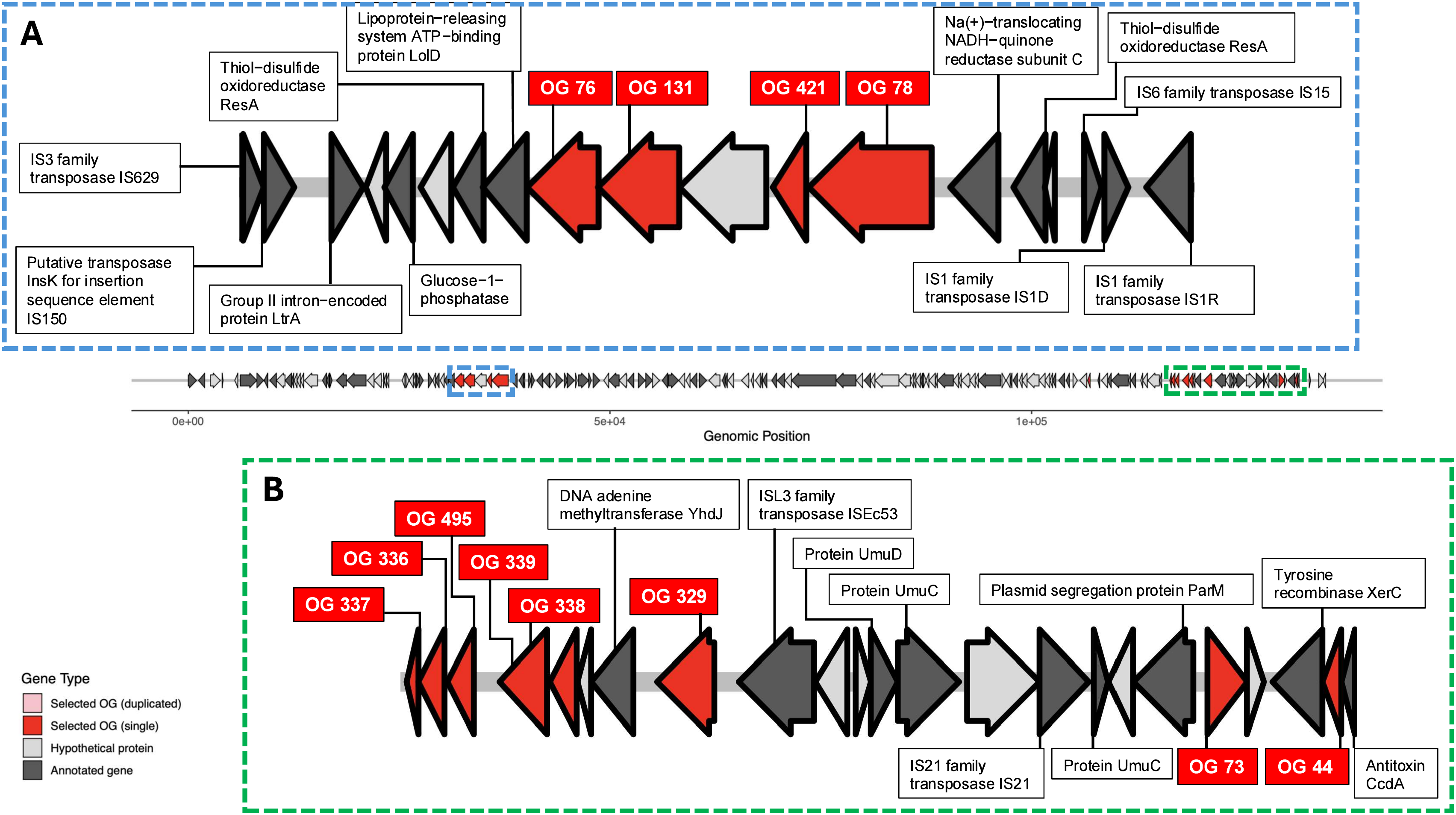
Location of the genes corresponding to the selected orthogroups (OGs) in the functional map of F plasmid pEIC35. (A) T region. (B) Y region.

**Figure 2.**
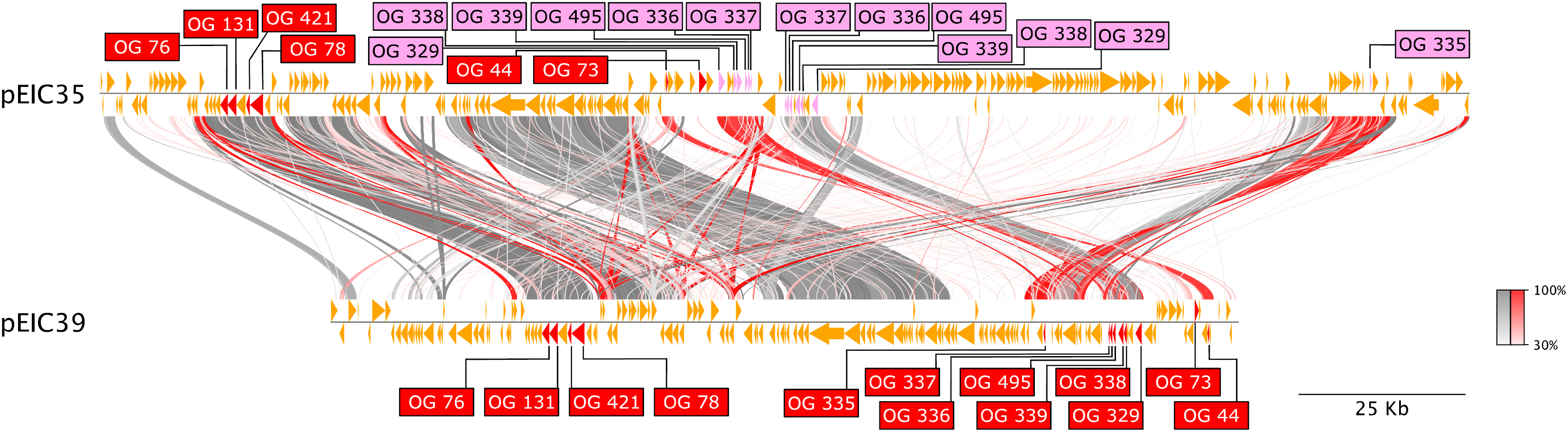
Location of the genes corresponding to the selected orthogroups (OGs) in the functional map of F plasmid pEIC39. (A) T region. (B) Y region.

Region T is flanked by transposable elements and contains numerous conserved genes, including the four OGs investigated in this study. The organization of this region is highly similar in pEIC35 and pEIC39 (Figs. 1A and 2A). In addition to OGs 76, 78, 131 and 421, the region contains genes encoding several proteins associated with metabolic or transport functions, including glucose-1-phosphatase, thiol-disulfide oxidoreductase (ResA), and the lipoprotein-releasing system ATP-binding protein LolD. According to the OG assignments, glucose-1-phosphatase and thiol-disulfide oxidoreductase (ResA) correspond to OGs 105 and 132, respectively. Both OGs were present in 79% of the F plasmids analyzed, just below the 83% threshold used to define the highly prevalent OGs in our initial analysis. LolD corresponds to OG296 and was detected in only 10% of the F plasmids analyzed.

The region Y also contains several genes involved in DNA metabolism. In contrast to region T, the region Y is not completely conserved in F plasmids pEIC35 and pEIC39. The genes corresponding to OGs 329 (encoding the DNA adenine methylase YhdJ), 336, 337, 338, and 339 are clustered together and occur in an inverted orientation within a conserved segment of both plasmids (Figs. 1B and 2B). The region also contains frequent insertion sequences, suggesting that this cluster may have been acquired or maintained as a single genetic unit. Region Y also contains several genes involved in DNA metabolism, including *xerC,* encoding a putative site-specific recombinase/resolvase, and *ltrA*, encoding the putative methyltransferase YubD. They correspond to highly prevalent OGs (OG498 and OG497, detected in 86% and 93% of the F plasmids analyzed, respectively). These OGs were not included in our analysis because proteins associated with DNA metabolism were excluded from the selection criteria. In pEIC35, several of the OGs identified in region Y are present as duplicates.

Importantly, the comparison of the two plasmids pEIC35 and pEIC39 (Fig. 3) showed that regions T and Y are among the three most highly conserved regions shared by the two plasmids, the third being primarily composed of *tra* genes (Supplementary Table S14). Thus, despite differences in the overall organization and conservation of region Y, both regions contain clusters of genes that are conserved between these two ST131 F plasmids.

**Figure 3.**
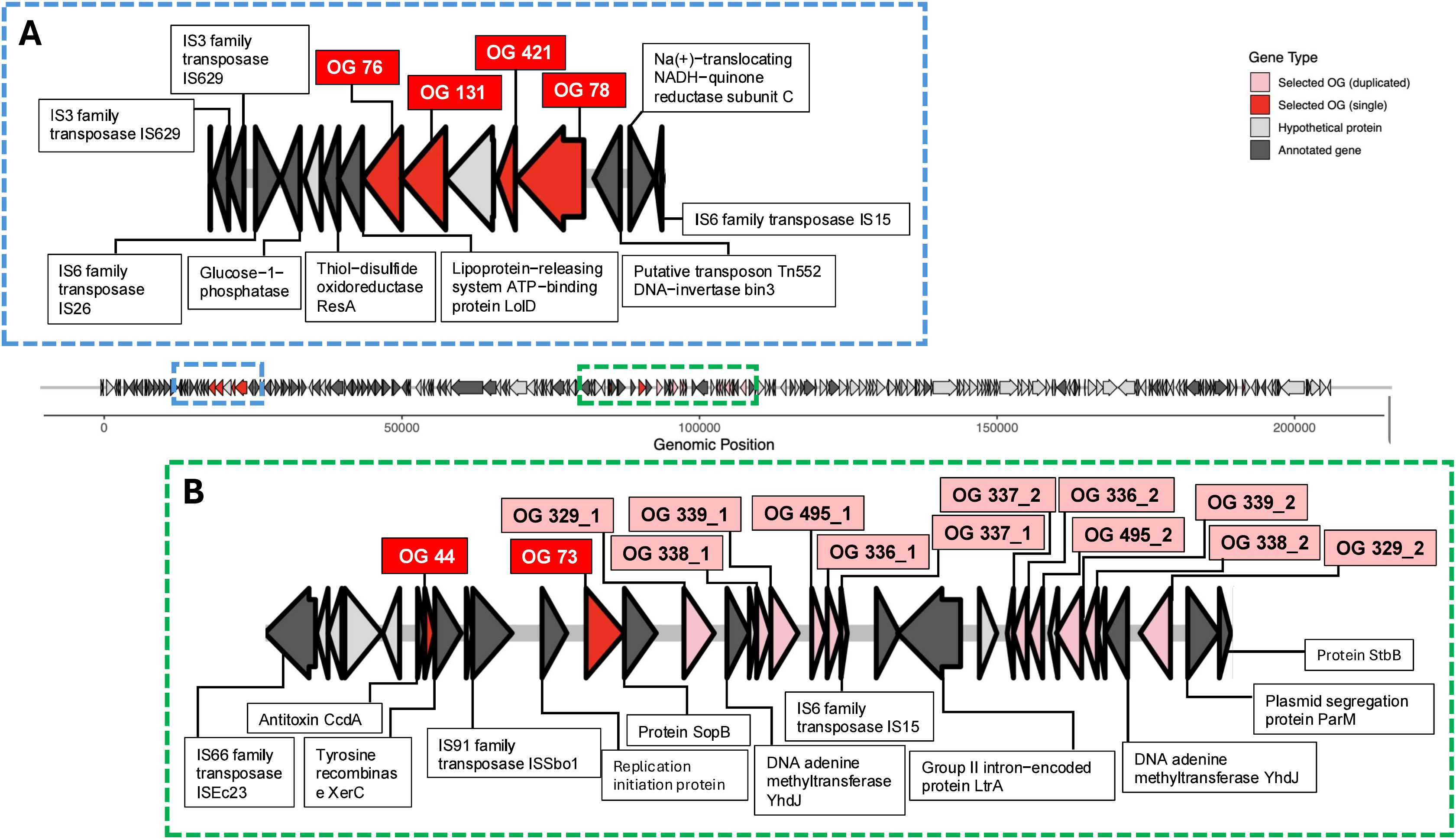
Comparative genomic analysis of the complete pEIC35 and pEIC39 F plasmids sequences based on protein sequence similarities.

## Discussion

Searching for conserved genes in the F plasmids of a group of *E. coli* ST131 isolates revealed additional features of F-plasmid genomics. Despite the considerable genomic diversity of F plasmids (23, 29, 30, 51), those present in all 29 ST131 strains contained several ORFs that were shared by more than 80% of the plasmids analyzed. These ORFs corresponded to either core or to cargo gene functions. Our subsequent analysis focused primarily on conserved ORFs of unknown or predicted function, which are likely to correspond to cargo genes. Two additional ORFs corresponding to proteins with known functions associated with plasmid DNA maintenance –SopA, involved in the stable distribution of plasmid copies, and CcdB together with CcdA, involved in plasmid stability– were included as controls. Based on their predicted function, when known, and their genomic locations, these OGs could be clearly divided into two groups: one comprising the ORFs in OGs 78,131, 76 and 421 (group T), and the other comprising the ORFs in OGs 329, 335, 336, 337, 338 and 339 (group Y).

Analysis of the presence of the different selected OGs in the *E. coli* F plasmids included in the COMPASS database (47) showed that only a small percentage of the plasmids from the sequenced strains –likely representing extraintestinal pathogenic *E. coli*– carried genes from both group Y and group T. In contrast, approximately 70% of the F plasmids harbored most, if not all, of the OGs belonging to group Y, indicating that the genes encoding these proteins are broadly distributed among F plasmids. Notably, the two YubG and the two YubJ OGs were co-inherited in most strains in the COMPASS collection. This pattern suggests that their gene products may have related or complementary roles in F-plasmid biology, and that the YubG and YubJ proteins within each pair may represent paralogues.

The genes encoding group T OGs clustered with other F-plasmid genes encoding proteins involved in metabolic and transport-related functions, suggesting that this region may contribute to the fitness of *E. coli* strains harboring these plasmids. These genes were particularly frequent in F plasmids from extraintestinal *E. coli* isolates and were highly represented among ST131 strains. In contrast, group Y OGs were located alongside genes associated with F-plasmid DNA metabolism, suggesting that these proteins of unknown function may participate in plasmid DNA-related processes.

A relevant aspect of this study is the presence of the selected OGs in the F plasmids of commensal *E. coli* strains, which have been considered important reservoirs of antibiotic resistance determinants in these strains (23). When individual plasmids of these strains were examined, the prevalence of group T or group Y OGs was clearly lower than that observed in plasmids from pathogenic *E. coli* strains (Table 4). However, when the entire plasmidome of each strain was considered, the prevalence of the identified OGs increased substantially (Table 4). Thus, individual commensal *E. coli* strains often collectively carry most of the OGs identified in this study. This observation raises the possibility that recombination among resident plasmids within a given strain could generate plasmids carrying combinations of most of the determinants characteristic of F plasmids from extraintestinal *E. coli* pathogens. Consequently, the plasmidome of commensal *E. coli* strains may represent not only a reservoir of antibiotic resistance genes, but also of fitness-associated genes that could facilitate bacterial colonization and survival in extraintestinal environments, as previously suggested (48, 52–54). This hypothesis warrants further investigation.

Mapping each OG onto the publicly available functionally annotated sequences of two F plasmids from two distinct ST131 strains provided important insights into the organization of the regions where these OGs map. The two plasmids differ considerably in size: the F plasmid from strain LREC 291 is substantially shorter (135,858 bp, 157 ORFs) than that from LREC 268 (205,049 bp, 228 ORFs). Despite these differences, Regions T and Y, which contain the two groups of OGs identified in this study, are highly similar in both plasmids, suggesting that these regions, together with additional genes, may be inherited as conserved genetic units. In both plasmids, genes encoding transposases are located both within and flanking these regions, consistent with their association with mobile genetic elements. This organization may explain why most strains carrying group T OGs contain all four genes, whereas most strains carrying group Y OGs harbor most of the genes belonging to this group.

Both F-plasmid regions contain genes that may contribute to the fitness and persistence of pathogenic *E. coli*: region T appears to be particularly associated with extraintestinal strains, whereas region Y is more broadly distributed among pathogenic *E. coli* strains. Region T, which also contains other conserved OGs such as OG132 (*resA*) and OG105 (glucose-1-phosphatase), forms a conserved metabolic cluster that is predominantly found in the F plasmids from extraintestinal *E. coli* strains (approximately 50%) and is particularly prevalent among ST131 strains (approximately 80%). Region Y, in addition to the Yub genes encoding proteins of unknown function, contains several genes associated with DNA metabolism. Notably, this region is duplicated in the pEIC35 plasmid, further supporting the possibility that these genes play an important role in F-plasmid biology (Fig. 1B). Given the known functions of several other genes within Region Y, the proteins encoded by the OGs identified in this study may also be involved in F-plasmid DNA-related processes. It is therefore intriguing that, despite the widespread distribution of these OGs among F plasmids, their functions have remained elusive.

The wide dissemination and potential relevance of OGs in regions T and Y are further supported by comparison with the two plasmid-maintenance genes included as controls, *sopA* and *ccdB*. Across the different F-plasmid collections analyzed, the prevalence of the OGs corresponding to these genes was never higher than that of the OGs from Regions T and Y (Tables 2-4). This was particularly evident in the OGs from Region T in extraintestinal *E. coli* isolates, especially ST131 strains, and for those from Region Y in pathogenic *E. coli* isolates.

Our study has identified two highly prevalent genomic regions in F plasmids that may contribute to the biology and fitness of their *E. coli* hosts. In particular, the genes within Region T may play relevant roles in the fitness of extraintestinal *E. coli*, specifically ST131 strains. This finding may have several potential applications, ranging from the identification of novel markers for the detection of high-risk clones such as ST131 in clinical isolates, to the identification of potential targets for limiting the dissemination of F plasmids among extraintestinal *E. coli* isolates or interfering with infections caused by MDR strains carrying these plasmids.

For the OGs encoded in Region Y, determining the functions of these conserved proteins could provide new insights into F-plasmid biology. Given their widespread distribution and their association with other genes involved in plasmid DNA-related processes, functional characterization of these proteins may also identify novel targets for strategies aimed at limiting the maintenance or dissemination of F plasmids carrying antibiotic resistance determinants in MDR *E. coli*.

## Acknowledgments

We thank María de Toro (Plataforma de Genómica y Bioinformática, Centro de Investigación Biomédica de La Rioja (CIBIR), Logroño, Spain) for her help with genome sequencing, library construction, and assembly.

## References

1. World Health Organization. 10 global health issues to track in 2021. Geneva: WHO; 2020. Accessed 21 Oct 2026.

2. Morens DM, Folkers GK, Fauci AS. The challenge of emerging and re-emerging infectious diseases. Nature. 2004;430:242–249. doi:10.1038/nature02759.

3. Antimicrobial Resistance Collaborators. Global burden of bacterial antimicrobial resistance in 2019: a systematic analysis. Lancet. 2022;399(10325):629–655. doi:10.1016/S0140-6736(21)02724-0.

4. GBD 2021 Antimicrobial Resistance Collaborators. Global burden of bacterial antimicrobial resistance 1990–2021: a systematic analysis with forecasts to 2050. Lancet. 2024;404(10459):1199–1226. doi:10.1016/S0140-6736(24)01867-1.

5. Carattoli A. Resistance plasmid families in Enterobacteriaceae. Antimicrob Agents Chemother. 2009;53(6):2227–2238. doi:10.1128/AAC.01707-08.

6. Carattoli A. Plasmids and the spread of resistance. Int J Med Microbiol. 2013;303(6–7):298–304. doi:10.1016/j.ijmm.2013.02.001.

7. Thomas CM, Nielsen KM. Mechanisms of, and barriers to, horizontal gene transfer between bacteria. Nat Rev Microbiol. 2005;3(9):711–721. doi:10.1038/nrmicro1234.

8. Wang J, Stephan R, Zurfluh K, Hächler H, Fanning S. Characterization of the genetic environment of blaESBL genes, integrons and toxin-antitoxin systems identified on large transferrable plasmids in multi-drug resistant *Escherichia coli*. Front Microbiol. 2015;5:716. doi:10.3389/fmicb.2014.00716.

9. Vrancianu CO, Popa LI, Bleotu C, Chifiriuc MC. Targeting plasmids to limit acquisition and transmission of antimicrobial resistance. Front Microbiol. 2020;11:761. doi:10.3389/fmicb.2020.00761.

10. Novick RP, Richmond MH. Nature and interactions of the genetic elements governing penicillinase synthesis in *Staphylococcus aureus*. J Bacteriol. 1965;90(2):467–480. doi:10.1128/jb.90.2.467-480.1965.

11. Hamad B. The antibiotics market. Nat Rev Drug Discov. 2010;9(9):675–676. doi:10.1038/nrd3267.

12. Baquero F, Coque TM, de la Cruz F. Ecology and evolution as targets: the need for novel eco-evo drugs and strategies to fight antibiotic resistance. Antimicrob Agents Chemother. 2011;55(8):3649–3660. doi:10.1128/AAC.00013-11.

13. Lipsitch M, Siber GR. How can vaccines contribute to solving the antimicrobial resistance problem? mBio. 2016;7(3):e00428–16. doi:10.1128/mBio.00428-16.

14. Lopatkin AJ, Meredith HR, Srimani JK, Pfeiffer C, Durrett R, You L. Persistence and reversal of plasmid-mediated antibiotic resistance. Nat Commun. 2017;8:1689. doi:10.1038/s41467-017-01532-1.

15. Hüttener M, Prieto A, Aznar S, Bernabeu M, Glaria E, Valledor AF, et al. Expression of a novel class of bacterial Ig-like proteins is required for IncHI plasmid conjugation. PLoS Genet. 2019;15(9):e1008399. doi:10.1371/journal.pgen.1008399.

16. Hüttener M, Hergueta J, Bernabeu M, Prieto A, Aznar S, Merino S, et al. Roles of proteins containing immunoglobulin-like domains in the conjugation of bacterial plasmids. mSphere. 2022;7(1):e00978–21. doi:10.1128/msphere.00978-21.

17. Prieto A, Miró L, Margolles Y, Bernabeu M, Salguero D, Merino S, et al. Targeting plasmid-encoded proteins that contain immunoglobulin-like domains to combat antimicrobial resistance. eLife. 2024;13:RP95328. doi:10.7554/eLife.95328.

18. Holt KE, Phan MD, Baker S, Duy PT, Nga TVT, Nair S, et al. Emergence of a globally dominant IncHI1 plasmid type associated with multiple drug resistant typhoid. PLoS Negl Trop Dis. 2011;5(7):e1245. doi:10.1371/journal.pntd.0001245.

19. Rozwandowicz M, Brouwer MSM, Fischer J, Wagenaar JA, Gonzalez-Zorn B, Guerra B, et al. Plasmids carrying antimicrobial resistance genes in Enterobacteriaceae. J Antimicrob Chemother. 2018;73(5):1121–1137. doi:10.1093/jac/dkx488.

20. Chong Y, Shimoda S, Shimono N. Current epidemiology, genetic evolution and clinical impact of extended-spectrum β-lactamase-producing *Escherichia coli* and *Klebsiella pneumoniae*. Infect Genet Evol. 2018;61:185–188. doi:10.1016/j.meegid.2018.04.005.

21. Naseer U, Sundsfjord A. The CTX-M conundrum: dissemination of plasmids and *Escherichia coli* clones. Microb Drug Resist. 2011;17(1):83–97. doi:10.1089/mdr.2010.0132.

22. Nicolas-Chanoine MH, Bertrand X, Madec JY. *\*Escherichia coli\** ST131, an intriguing clonal group. Clin Microbiol Rev. 2014;27(3):543–574. doi:10.1128/CMR.00125-13.

23. Stephens C, Arismendi T, Wright M, Hartman A, Gonzalez A, Gill M, et al. F plasmids are the major carriers of antibiotic resistance genes in human-associated commensal *Escherichia coli*. mSphere. 2020;5(4):e00709–20. doi:10.1128/mSphere.00709-20.

24. Koraimann G. Spread and persistence of virulence and antibiotic resistance genes: a ride on the F plasmid conjugation module. EcoSal Plus. 2018;8(1). doi:10.1128/ecosalplus.ESP-0003-2018.

25. Haiko J, Suomalainen M, Ojala T, Lähteenmäki K, Korhonen TK. Invited review: breaking barriers—attack on innate immune defences by omptin surface proteases of enterobacterial pathogens. Innate Immun. 2009;15(2):67–80. doi:10.1177/1753425909102559.

26. Troxell B, Hassan HM. Transcriptional regulation by Ferric Uptake Regulator (Fur) in pathogenic bacteria. Front Cell Infect Microbiol. 2013;3:59. doi:10.3389/fcimb.2013.00059.

27. Fleckenstein JM, Hardwidge PR, Munson GP, Rasko DA, Sommerfelt H, Steinsland H. Molecular mechanisms of enterotoxigenic *Escherichia coli* infection. Microbes Infect. 2010;12(2):89–98. doi:10.1016/j.micinf.2009.10.002.

28. Cusumano CK, Hung CS, Chen SL, Hultgren SJ. Virulence plasmid harbored by uropathogenic *Escherichia coli* functions in acute stages of pathogenesis. Infect Immun. 2010;78(4):1457–1467. doi:10.1128/IAI.01260-09.

29. Lanza VF, de Toro M, Garcillán-Barcia MP, Mora A, Blanco J, Coque TM, et al. Plasmid flux in *Escherichia coli* ST131 sublineages, analyzed by plasmid constellation network (PLACNET), a new method for plasmid reconstruction from whole genome sequences. PLoS Genet. 2014;10(12):e1004766. doi:10.1371/journal.pgen.1004766.

30. Wyrsch ER, Bushell RN, Marenda MS, Browning GF, Djordjevic SP. Global phylogeny and F virulence plasmid carriage in pandemic *Escherichia coli* ST1193. Microbiol Spectr. 2022;10(6):e02554–22. doi:10.1128/spectrum.02554-22.

31. Flament-Simon SC, Nicolas-Chanoine MH, García V, Duprilot M, Mayer N, Alonso MP, García-Meniño I, et al. Clonal structure, virulence factor-encoding genes and antibiotic resistance of *Escherichia coli* causing urinary tract infections and other extraintestinal infections in humans in Spain and France during 2016. Antibiotics (Basel). 2020;9(4):161. doi:10.3390/antibiotics9040161.

32. Price LB, Johnson JR, Aziz M, Clabots C, Johnston B, Tchesnokova V, et al. The epidemic of extended-spectrum-β-lactamase-producing *Escherichia coli* ST131 is driven by a single highly pathogenic subclone, H30-Rx. mBio. 2013;4(6):e00377-13. doi:10.1128/mBio.00377-13.

33. Johnson JR, Clermont O, Johnston B, Clabots C, Tchesnokova V, Sokurenko E, et al. Rapid and specific detection, molecular epidemiology, and experimental virulence of the O16 subgroup within *Escherichia coli* sequence type 131. J Clin Microbiol. 2014;52(5):1358–1365. doi:10.1128/JCM.03502-13.

34. Blanc V, Leflon-Guibout V, Blanco J, Haenni M, Madec JY, Rafignon G, et al. Prevalence of day-care centre children (France) with faecal CTX-M-producing *Escherichia coli* comprising O25b:H4 and O16:H5 ST131 strains. J Antimicrob Chemother. 2014;69(5):1231–1237. doi:10.1093/jac/dkt519.

35. Petty NK, Ben Zakour NL, Stanton-Cook M, Skippington E, Totsika M, Forde BM, et al. Global dissemination of a multidrug resistant *Escherichia coli* clone. Proc Natl Acad Sci U S A. 2014;111(15):5694–5699. doi:10.1073/pnas.1322678111.

36. Coque TM, Novais A, Carattoli A, Poirel L, Pitout J, Peixe L, et al. Dissemination of clonally related *Escherichia coli* strains expressing extended-spectrum β-lactamase CTX-M-15. Emerg Infect Dis. 2008;14(2):195–200. doi:10.3201/eid1402.070350.

37. Pitout JDD, Finn TJ. The evolutionary puzzle of *Escherichia coli* ST131. Infect Genet Evol. 2020;81:104265. doi:10.1016/j.meegid.2020.104265.

38. Flament-Simon SC, García V, Duprilot M, Mayer N, Alonso MP, García-Meniño I, et al. High prevalence of ST131 subclades C2-H30Rx and C1-M27 among extended-spectrum β-lactamase-producing *Escherichia col*i causing human extraintestinal infections in patients from two hospitals of Spain and France during 2015. Front Cell Infect Microbiol. 2020;10:125. doi:10.3389/fcimb.2020.00125.

39. Vielva L, de Toro M, Lanza VF, de la Cruz F. PLACNETw: a web-based tool for plasmid reconstruction from bacterial genomes. Bioinformatics. 2017;33(23):3796–3798. doi:10.1093/bioinformatics/btx462.

40. Seemann T. Prokka: rapid prokaryotic genome annotation. Bioinformatics. 2014;30(14):2068–2069. doi:10.1093/bioinformatics/btu153.

41. Derelle R, Philippe H, Colbourne JK. Broccoli: combining phylogenetic and network analyses for orthology assignment. Mol Biol Evol. 2020;37(11):3389–3396. doi:10.1093/molbev/msaa159.

42. Altschul SF, Gish W, Miller W, Myers EW, Lipman DJ. Basic local alignment search tool. J Mol Biol. 1990;215(3):403–410. doi:10.1016/S0022-2836(05)80360-2.

43. McGinnis S, Madden TL. BLAST: at the core of a powerful and diverse set of sequence analysis tools. Nucleic Acids Res. 2004;32(Web Server issue):W20–W25. doi:10.1093/nar/gkh435.

44. Huerta-Cepas J, Forslund K, Coelho LP, Szklarczyk D, Jensen LJ, von Mering C, et al. Fast genome-wide functional annotation through orthology assignment by eggNOG-mapper. Mol Biol Evol. 2017;34(8):2115–2122. doi:10.1093/molbev/msx148.

45. Cantalapiedra CP, Hernández-Plaza A, Letunic I, Bork P, Huerta-Cepas J. eggNOG-mapper v2: functional annotation, orthology assignments, and domain prediction at the metagenomic scale. Mol Biol Evol. 2021;38(12):5825–5829. doi:10.1093/molbev/msab293.

46. Jones P, Binns D, Chang HY, Fraser M, Li W, McAnulla C, et al. InterProScan 5: genome-scale protein function classification. Bioinformatics. 2014;30(9):1236–1240. doi:10.1093/bioinformatics/btu031.

47. Douarre PE, Mallet L, Radomski N, Felten A, Mistou MY. Analysis of COMPASS, a new comprehensive plasmid database revealed prevalence of multireplicon and extensive diversity of IncF plasmids. Front Microbiol. 2020;11:483. doi:10.3389/fmicb.2020.00483.

48. Reid CJ, Cummins ML, Djordjevic SP. Major F plasmid clusters are linked with ColV and pUTI89-like marker genes in bloodstream isolates of *Escherichia coli*. BMC Genomics 2025;26:57. doi:10.1186/s12864-025-11226-4.

49. Cock PJA, Antao T, Chang JT, Chapman BA, Cox CJ, Dalke A, et al. Biopython: freely available Python tools for computational molecular biology and bioinformatics. Bioinformatics. 2009;25(11):1422–1423. doi:10.1093/bioinformatics/btp163.

50. Shimoyama Y. pyGenomeViz: a genome visualization Python package for comparative genomics [software]. Released 18 May 2024. Accessed 21 Oct 2026.

51. Al Mamun AAM, Kissoon K, Li YG, Hancock E, Christie PJ. The F plasmid conjutome: the repertoire of *E. coli* proteins translocated through an F-encoded type IV secretion system. mSphere. 2024;9(7):e0035424. doi:10.1128/msphere.00354-24.

52. Le Gall T, Clermont O, Gouriou S, Picard B, Nassif X, Denamur E, et al. Extraintestinal virulence is a coincidental by-product of commensalism in B2 phylogenetic group *Escherichia coli* strains. Mol Biol Evol. 2007;24(11):2373–2384. doi:10.1093/molbev/msm172.

53. Östblom A, Adlerberth I, Wold AE, Nowrouzian FL. Pathogenicity island markers, virulence determinants malX and usp, and the capacity of *Escherichia coli* to persist in infants’ commensal microbiotas. Appl Environ Microbiol. 2011;77(7):2303–2308. doi:10.1128/AEM.02405-10.

54. Thänert R, Reske KA, Hink T, Wallace MA, Wang B, Schwartz DJ, et al. Comparative genomics of antibiotic-resistant uropathogens implicates three routes for recurrence of urinary tract infections. mBio. 2019;10(4):e01977–19. doi:10.1128/mBio.01977-19.

