## Supplementary methods for "Highly prevalent cargo genes of F plasmids harbored in pathogenic *Escherichia coli*"

**SUPPLEMENTARY MATERIALS AND METHODS**

**In silico typing and characterization of *E. coli* isolates**

Genome assemblies were characterized *in silico* using tools available through the Center for Genomic Epidemiology (CGE) platform and complementary databases. Phylogroups were assigned using Clermont Typing (1), sequence types (STs) were determined according to the Achtman MLST scheme, and O serotypes were identified using SerotypeFinder (Supplementary Table S15). Clone designation was based on the combination of phylogroup, clonotype (CH), and sequence type (ST), while ST131 isolates were further classified into their corresponding phylogenetic clades following the established population structure of this lineage (2).

Plasmid incompatibility groups and plasmid sequence types (pMLST) were determined using PlasmidFinder and pMLST, respectively (Supplementary Table S15). Virulence-associated genes were identified using VirulenceFinder and the Virulence Factor Database (VFDB) with minimum coverage and identity thresholds of 90% and 75%, respectively (Supplementary Table S15). Antimicrobial resistance genes, including extended-spectrum β-lactamase (ESBL)-encoding **bla**<sub>CTX-M</sub> genes, were identified using ResFinder, ARG-ANNOT, and the Comprehensive Antibiotic Resistance Database (CARD) with minimum coverage and identity thresholds of 93% and 75%, respectively (Supplementary Table S15). Point mutations associated with antimicrobial resistance were identified using PointFinder. Putative integrative and conjugative elements (ICEs) were identified using ICEberg (ICEfinder and VRprofile), and additional database searches were performed using MyDbFinder when required (Supplementary Table S15). CSI Phylogeny was used for whole-genome phylogenetic analysis.

**Plasmid reconstruction, typing and relaxase classification**

Plasmid reconstruction from WGS data was performed usingPLACNETw (3), a graph-based tool for plasmid reconstruction. However, manual pruning of the graphs is required, and not all plasmids could always be successfully separated without introducing some risk of error. For this reason, some plasmids remain merged and the resulting plasmidome reconstruction should be considered presumptive (Supplementary Tables S6 and S7). In the PLACNETw representation, most plasmids and integrative conjugative elements (ICEs) can be recognized by their replication initiation proteins (RIP) and/or RELaxase proteins (REL), which are considered as markers of Mobile Genetic Elements (MGEs).

Incompatibility groups (Inc) and pMLST subtypes were *in silico* determined using PlasmidFinder and pMLST (4), respectively, via the CGE online services (<http://genomicepidemiology.org/>; Supplementary Table S15). Additionally, reconstructed plasmids were subtyped according to their relaxase family byphylogenetic comparison with previously defined relaxase subfamilies (5) (Supplementary Tables S6, S7 and S15).

Transferability was presumptively deduced based on the identification of relaxases (mobilizable plasmids) and mating pair formation (Mpf) system proteins (conjugative plasmids).

The reconstructed plasmidome was described according to plasmid size, estimated copy number, relaxase family (MOB), incompatibility group (Inc.), and the associated virulence and antimicrobial resistance genes. Approximate plasmid-copy number was estimated from the ratio between plasmid and chromosome sequencing coverage as determined by PLACNETw (Supplementary Tables S6 and S7).

**References Supplementary materials and methods**:

1. Clermont O, Christenson JK, Denamur E, Gordon DM. The Clermont *Escherichia coli* phylo-typing method revisited: improvement of specificity and detection of new phylo-groups. **Environ Microbiol Rep**. 2013;5(1):58–65. doi:10.1111/1758-2229.12019.

2. Pitout JDD, Finn TJ. The evolutionary puzzle of *Escherichia coli* ST131. **Infect Genet Evol.** 2020;81:104265. doi:10.1016/j.meegid.2020.104265.

3. Vielva L, de Toro M, Lanza VF, de la Cruz F. PLACNETw: a web-based tool for plasmid reconstruction from bacterial genomes. **Bioinformatics**. 2017;33(23):3796–3798. doi:10.1093/bioinformatics/btx462.

4. Carattoli A, Zankari E, García-Fernández A, Voldby Larsen M, Lund O, Villa L, et al. In silico detection and typing of plasmids using PlasmidFinder and plasmid multilocus sequence typing. **Antimicrob Agents Chemother**. 2014;58(7):3895–3903. doi:10.1128/AAC.02412-14.

5. Alvarado A, Garcillán-Barcia MP, de la Cruz F. A degenerate primer MOB typing (DPMT) method to classify gamma-proteobacterial plasmids in clinical and environmental settings. **PLoS One**. 2012;7(7):e40438. doi:10.1371/journal.pone.0040438.
